# Analyzing the differences in olfactory bulb mitral cell spiking with ortho- and retronasal stimulation

**DOI:** 10.1101/2021.06.14.448305

**Authors:** Michelle F. Craft, Andrea K. Barreiro, Shree Hari Gautam, Woodrow L. Shew, Cheng Ly

**Affiliations:** Department of Statistical Sciences and Operations Research, Virginia Commonwealth University, Richmond, VA 23284 U.S.A.; Department of Mathematics, Southern Methodist University, Dallas, TX 75275 U.S.A.; Department of Physics, University of Arkansas, Fayetteville, AR 72701 U.S.A.

## Abstract

The majority of olfaction studies focus on orthonasal stimulation where odors enter via the front nasal cavity, while retronasal olfaction, where odors enter the rear of the nasal cavity during feeding, is understudied. The processing of retronasal odors via coordinated spiking of neurons in the olfactory bulb (**OB**) is largely unknown. To this end, we use multi-electrode array *in vivo* recordings of rat OB mitral cells (**MC**) in response to a food odor with both modes of stimulation, and find significant differences in evoked firing rates and spike count covariances (i.e., noise correlations). To better understand these differences, we develop a single-compartment biophysical OB model that is able to reproduce key properties of important OB cell types. Prior experiments in olfactory receptor neurons (**ORN**) showed retro stimulation yields slower and spatially smaller ORN inputs than with ortho, yet whether this is consequential for OB activity remains unknown. Indeed with these specifications for ORN inputs, our OB model captures the trends in our OB data. We also analyze how first and second order ORN input statistics dynamically transfer to MC spiking statistics with a phenomenological linear-nonlinear filter model, and find that retro inputs result in larger temporal filters than ortho inputs. Finally, our models show that the temporal profile of ORN is crucial for capturing our data and is thus a distinguishing feature between ortho and retro stimulation, even at the OB. Using data-driven modeling, we detail how ORN inputs result in differences in OB dynamics and MC spiking statistics. These differences may ultimately shape how ortho and retro odors are coded.

**Author summary:** Olfaction is a key sense for many cognitive and behavioral tasks, and is particularly unique because odors can naturally enter the nasal cavity from the front or rear, i.e., ortho- and retro-nasal, respectively. Yet little is known about the differences in coordinated spiking in the olfactory bulb with ortho versus retro stimulation, let alone how these different modes of olfaction may alter coding of odors. We simultaneously record many cells in rat olfactory bulb to assess the differences in spiking statistics, and develop a biophysical olfactory bulb network model to study the reasons for these differences. Using theoretical and computational methods, we find that the olfactory bulb transfers input statistics differently for retro stimulation relative to ortho stimulation. Furthermore, our models show that the temporal profile of inputs is crucial for capturing our data and is thus a distinguishing feature between ortho and retro stimulation, even at the olfactory bulb. Understanding the spiking dynamics of the olfactory bulb with both ortho and retro stimulation is a key step for ultimately understanding how the brain codes odors with different modes of olfaction.

## Introduction

Olfactory processing naturally occurs in two distinct modes: orthonasal (**ortho**) where odors enter the front of the nasal cavity and retronasal (**retro**) where odors enter the rear. Despite the importance of retronasal olfaction, i.e., retro naturally occurs during feeding, it is relatively understudied. Specifically, it is unknown whether odors in higher brain regions are processed differently depending on the mode of olfaction (ortho versus retro). The olfactory bulb (**OB**) is the main area where odor information is processed and subsequently transferred to cortex via mitral cell (**MC**) (and tufted cell) spiking. The differences in MC spiking with ortho and retro stimulation have implications for odor processing, but any such differences are largely unknown.

Studies have shown that olfactory receptor neuron (**ORN**) activity, which are presynaptic to the OB, differ for ortho versus retro stimulation. This has been shown in various ways, including with fMRI [1], calcium imaging [2], and optical imaging in transgenic mice [3]. These and other prior studies [4–6] suggest that ORN inputs likely lead to any observed differences in OB activity. However, the implications of these differences in ORN for MC spiking have yet to be explored.

We perform *in vivo* recordings of rat OB mitral cells using multi-electrode arrays with a food odor stimulus, delivered by both modes of stimulation, to determine whether there are differences in MC population spiking. We find significant differences in odor-evoked MC spiking with ortho versus retro stimulation in both firing rate (larger with retro) and spike count covariance (larger with ortho). Dissecting how components of ORN inputs alter OB spiking is difficult experimentally due to the complexity of both the recurrent circuitry in the OB [7, 8] and resulting spatiotemporal ORN responses [4, 6]. So we developed a single-compartment biophysical OB model to investigate how ORN input statistics affect the first and second order MC spiking statistics. Specifically, we model ORN input as a time-varying inhomogeneous Poisson Process [9], where the input rate has slower increase and decay for retro than ortho [2, 3], and the ORN input correlation is smaller for retro than ortho [2, 3]. With these specifications, our biophysical OB network model is able to capture the ortho versus retro MC spiking response trends in our experimental data.

Understanding how retro stimulation can elicit both larger firing rates and smaller co-variablity than ortho is generally difficult in recurrent networks because of the numerous attributes that shape spike statistics [10–13]. Since our biophysical OB model is too complex to directly analyze, we use a linear-nonlinear (**LN**) model framework to analyze how our realistic OB network transfers input statistics (from ORN) to outputs (MC spike statistics). We find that with retro inputs, the OB network effectively filters input statistics (in time) with larger absolute values than with ortho inputs. Thus the OB network is more sensitive to fluctuations with retro-like inputs than with ortho. Finally, we use our models examine which of these attribute(s) of ORN inputs (temporal profile, amplitude, input correlation) are most significant for capturing our data. We find that temporal profile is the critical attribute for ortho versus retronasal stimulus response.

This work provides a framework for how to analyze the sources driving different OB spiking responses to different modes of olfaction, as well as important insights that have implications for how the brain codes odors.

## Results

We performed *in vivo* multi-electrode array recordings of the OB in the mitral cell layer of anesthetized rats (see **Materials and methods: Electrophysiological recordings**) to capture odor evoked spiking activity of populations of putative MCs. This yielded a large number of cells (94) and simultaneously recorded pairs of cells (1435) with which to assess population average spiking statistics. The first and second order spike statistics are summarized in Figure 1, including the firing rate (peri-stimulus time histogram, **PSTH**, Fig 1A), the spike count variance (Fig 1B), the spike count covariance (Fig 1C), Fano Factor (variance divided by mean, Fig 1D), and Pearson’s correlation (Fig 1E). Spike count statistics were calculated with half-overlapping 100 ms time windows. The time window 100 ms is an intermediate value between shorter (membrane time constants, AMPA, GABA_A_, etc.) and longer time scales (NMDA, calcium, and other ionic currents) known to exist in the OB.

**Fig 1.**
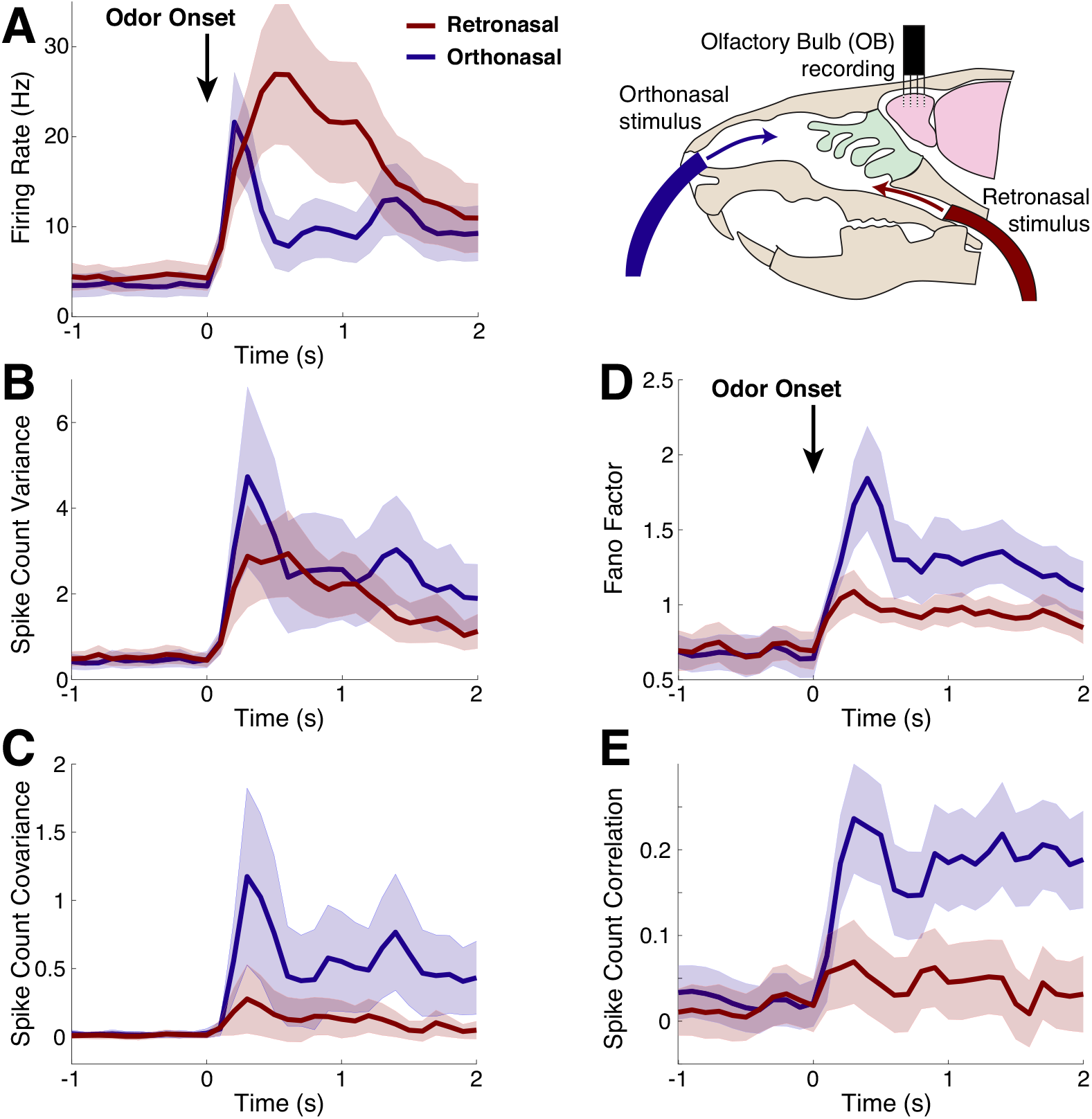
Spike statistics from *in vivo* multi-electrode array recordings. Population average spike statistics for orthonasal (blue) and retronasal (red) with stimulus onset at time *t* = 0 s as indicated by black arrow for 1 s duration. **A)** Firing rate (Hz) is statistically significantly different between ortho and retro for the duration of the evoked period (0.4 ≤ *t* ≤ 1.1 s, see Fig S1 in **S1. Supplementary Material**). **B)** Spike count variance has no statistical significant differences between ortho and retro. **C)** Covariance of spike counts are statistically significant different throughout the evoked state (0 ≤ *t* ≤ 2) with ortho having larger values. Scaled measures of variability shown for completeness: Fano Factor **(D)** is the variance divided by mean spike count, and Pearson’s correlation **(E)** is the covariance divided by the product of the standard deviations; both are also different with ortho versus retro. Spike counts in 100 ms half-overlapping time windows averaging over all 10 trials. Significance: two-sample t-tests (assuming unequal variances) for each time bin to assess differences in population means, *p* < 0.01. From 94 total cells and 1435 simultaneously recorded cell pairs; shaded regions show relative population heterogeneity: *μ* ± 0.2std (standard deviation across the population/pairs after trial-averaging; 0.2 scale for visualization).

We find statistically significant differences between ortho and retro stimulation in almost all of the first and second order MC spike count statistics. At odor onset, orthonasal stimulation elicited larger firing rates with a faster rise than retronasal, after which retronasal firing is larger and remains elevated longer than with orthonasal. These trends are consistent with imaging studies of the glomeruli layer in OB in transgenic mice (see [3], their figure 2) as well as EOG recordings of the superficial layers of the OB in rats (see [5], their figure 7). More specifically, we find statistical significance (*α* = 0.01) between ortho- and retronasal firing rate after and for the duration of the odor stimulation. We also find that MC spike count covariance for orthonasal stimulus was significantly larger than retronasal for the entirety of the evoked state. MC spike count variance, however, was not significantly different between ortho and retronasal stimulus. For detailed plots of significance in time of the first and second order spike statistics, see Fig S1 in **S1. Supplementary Material**. Hereafter, we mainly focus on understanding the differences in firing rate and spike count covariance because they directly impact common coding metrics (e.g. the Fisher information) in contrast to scaled measures of variability (Fano factor and Pearson’s correlation) which do not directly impact common coding metrics [14]. Moreover, Fano factor and correlation both depend on variance, which is not statistically different with ortho and retro (but see Fig S1D and S1E in **S1. Supplementary Material** for completeness).

**Fig 2.**
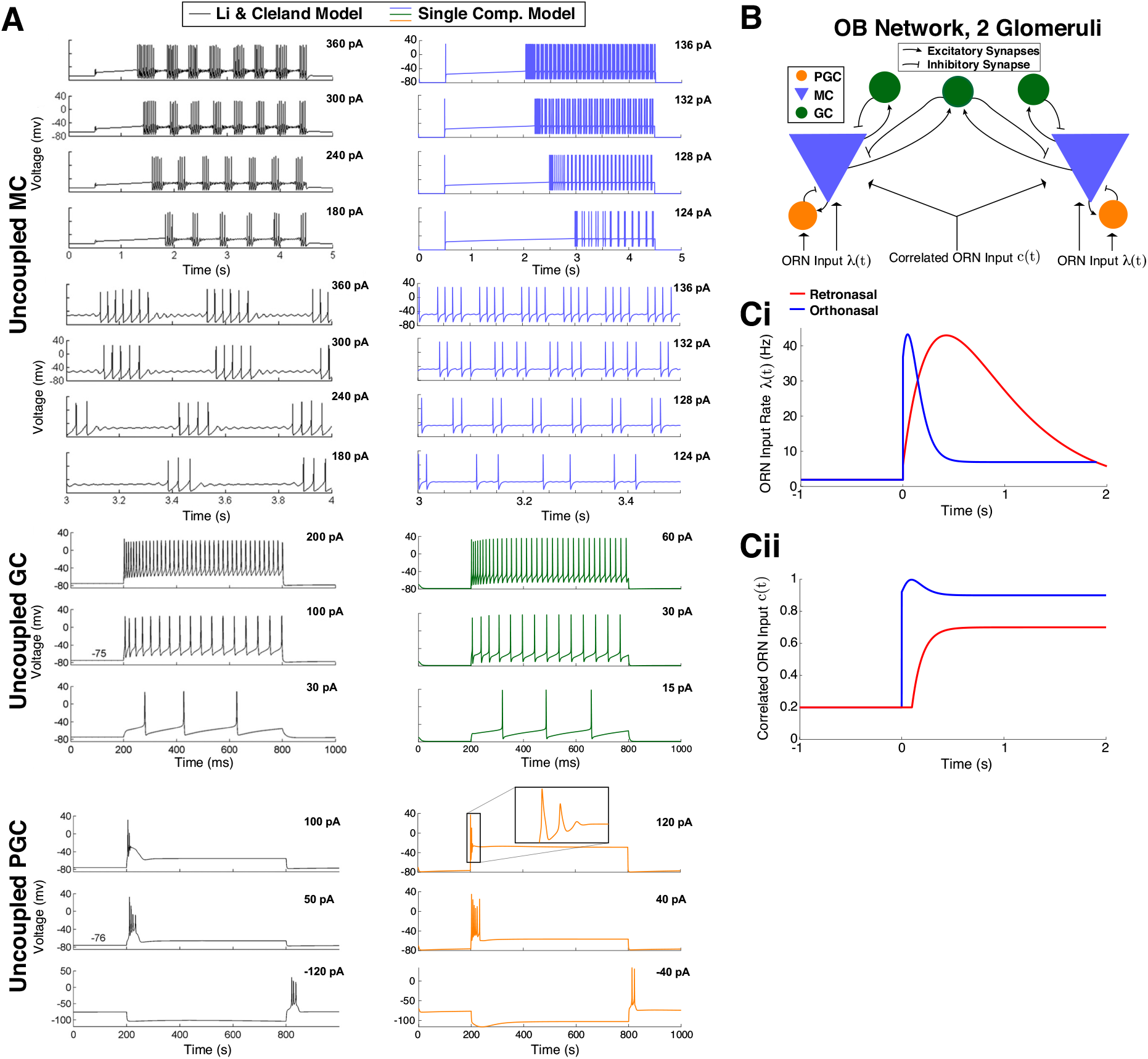
Biophysical OB model. **A)** Dynamics of the 3 uncoupled cell models. MC voltage dynamics with current step inputs in Li & Cleland models (black curves on the left, copied from Li & Cleland [15]) are captured by our single-compartment model (blue on the right). Rows 5–8 show expanded time view of first 4 rows to highlight spike cluster sizes and sub-threshold oscillations (same voltage axis for each). GC voltage responses to three different levels of current injection in the Li & Cleland model (black curves on the left) is similar to our model (green on the right). PGC responses with depolarizing current steps again are similar in both models. Note that release from a hyperpolarizing current injection leads to transient spiking in both models (bottom). **B)** Coupled OB network model of 2 glomeruli with ORN inputs. ORN synapses are driven by correlated inhomogeneous Poisson Processes (Eqs (9)–(11)). **C)**Based on ORN imaging studies, we set *λ_O_*(*t*) to increase and decay faster than *λ_R_*(*t*) with odor onset at time 0s (i). Similarly, we set the input correlation of ORN synapses to the 2 MCs to *c_R/O_*(*t*) where *c_R_*(*t*) *< c_O_*(*t*) and *c_O_*(*t*) rises quicker than *c_R_*(*t*) (ii).

### OB network model captures data trends

To better understand how differences in MC spiking with ortho and retro stimulation come about, we developed a single-compartment OB network model based on Li & Cleland’s multicompartment model [15, 16]. Our model is more computationally efficient than their larger multi-compartment models [15, 16], requiring a fraction of the variables (tens of state variables instead of thousands). Importantly, our single-compartment model retains important biophysical features (Fig 2A).

In Fig 2A, we see in both models of MC (uncoupled) that the time to spiking decreases with increasing current values, and the number of spikes in a cluster increases with current values consistent with prior electrophysiological experiments [17–19]. In our model, the spacing between spike clusters and number of spikes in a cluster in our model (right) qualitatively match the Li & Cleland model (left). The sub-threshold oscillations are not as prominent as in Li & Cleland, but still apparent. In the uncoupled GC models, both exhibit a delay to the first spike with weak current step [20] (Fig 2A, bottom) and tonic firing without appreciable delay for higher current injections [21] (Fig 2A, middle and top). In the uncoupled PGC models, we do not observe repetitive firing in either models (Fig 2A, top and middle). Also, releasing from a hyperpolarizing current injection (bottom) can illicit spiking in both models, as observed by McQuiston & Katz [22]. Thus, we have condensed the OB model by using far less equations than Cleland’s models while retaining many of the biophysical dynamics known to exist in these 3 important OB cell types.

Since our focus is on first and second order population-averaged spiking statistics, we use a minimal OB network model with 2 glomeruli (Fig. 2B). Each glomerulus has a PGC, MC and GC cell; we also include a common GC that provides shared inhibition to both MCs because GCs are known to span multiple glomeruli and shape MC spike correlation [8, 23, 24]. Within the OB network, the PGC and GC cells provide presynaptic GABA_A_ inhibition to the MCs they are coupled to, while MC provide both AMPA and NMDA excitation to PGC and GCs (see **Materials and methods: Single-Compartment Biophysical Model** for further details). The ORN synaptic inputs are an important component of this coupled OB network; they are driven by correlated inhomogeneous Poisson Process with increases in rate and correlation at odor onset. The specific time-varying input rate and correlation we use are shown in Figs. 2Ci and Cii, respectively. The differences in ortho versus retro (Fig. 2Ci and Cii) are based on prior studies of ORN input to the OB in response to both ortho and retro stimulation [2, 3].

A comparison of first and second-order statistics between our OB model and *in vivo* data is shown in Fig 3. With the ORN activity specified in Fig 2C, our OB model is able to qualitatively capture trends seen in our data. Firing rates in Fig 3A show that both the model and data exhibit larger firing rates for ortho at odor onset followed by a sharper decline. After the initial increase in ortho firing rates, retro firing rates continue to increase, eventually becoming larger than ortho and remaining elevated longer, consistent with optical imaging experiments (see [3] their Fig 2). Although there is no significant differences in the spike count variance between ortho and retro in our experimental data, we show our data with model for completeness (Fig 3B).

**Fig 3.**
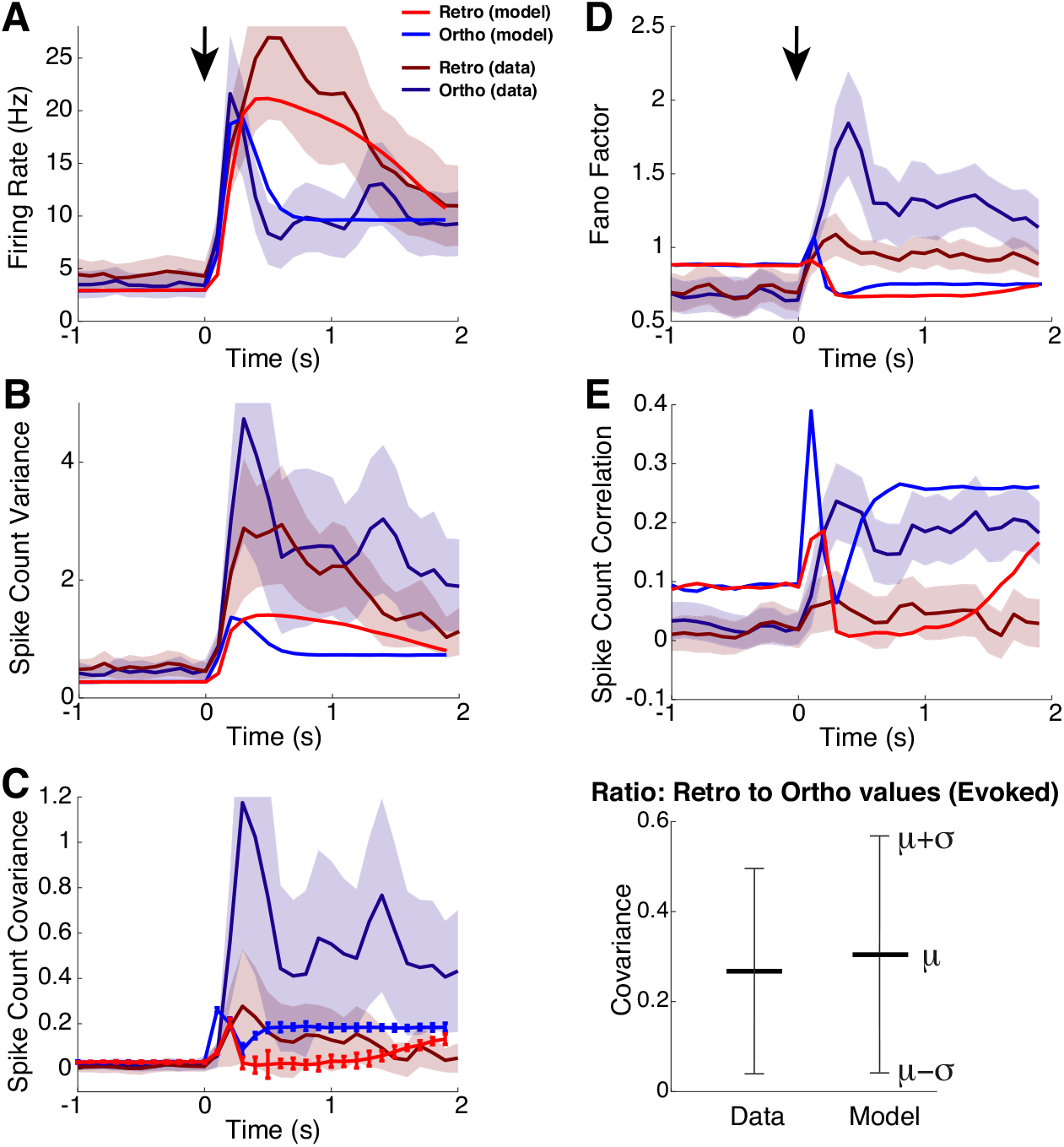
OB model captures trends in our data. Comparison of all first and second-order statistics of coupled OB network model to our data. **A)** Firing rate of ortho increases and decays faster than retro in both data and model. **B)** Variance of spike counts for ortho and retro shown for completeness, but recall that in experimental data that they are not statistically different. **C)** Covariance of spike counts is larger for ortho than retro in both data and model (left), but the magnitudes of data and model differ. Right: comparison of the ratio of retro covariance to ortho covariance in the evoked state, *t* ≥ 0 s, shows that the model captures the relative differences between ortho and retro. Comparisons of the (**D**) Fano factor and (**E**) Pearson’s spike count correlation shown for completeness despite both measures depending on spike count variance. **D)** The model has slightly larger Fano factor with ortho, consistent with the data. **E)** The model does qualitatively capture the spike count correlation for both ortho and retro, at least in the evoked state.

Our OB model captures the trend that ortho spike count covariance is larger than retro after odor onset, Fig 3C (left). The OB model certainly does not capture the magnitude of the spike count covariance in the data; recall that covariance in our experimental data is the population average over all 1435 simultaneously recorded pairs with significant heterogeneity while our model is homogeneous. But the relative differences between retro and ortho (as measured by the ratio of retro to ortho covariance in the evoked state) are similar (Fig 3C, right). Thus our OB model captures the trends of the population-averaged spike count statistics. We also show comparisons of Fano Factor (Fig 3D) and Pearson’s correlation (Fig 3E) for completeness. Consistent with our data, our OB model has larger Fano Factor and spike count correlation for ortho than with retro. In the evoked state, the OB model matches spike count correlation for both ortho and retro well. The larger ortho Fano factor in our data is captured in our model, but the difference is very modest.

### How OB network transfers ORN input statistics

We next sought to better understand how our OB network model operates with different ORN inputs. In particular, we investigated whether the same OB network model transferred ortho and retro ORN inputs to MC spike outputs differently or not. We addressed this in a simple and transparent manner, using a phenomenological LN model (Fig 4A) to approximate the overall effects of the OB network on ORN inputs. LN-type models have often been to circumvent the complexities in biophysical spiking models (see [25–27] and Discussion).

**Fig 4.**
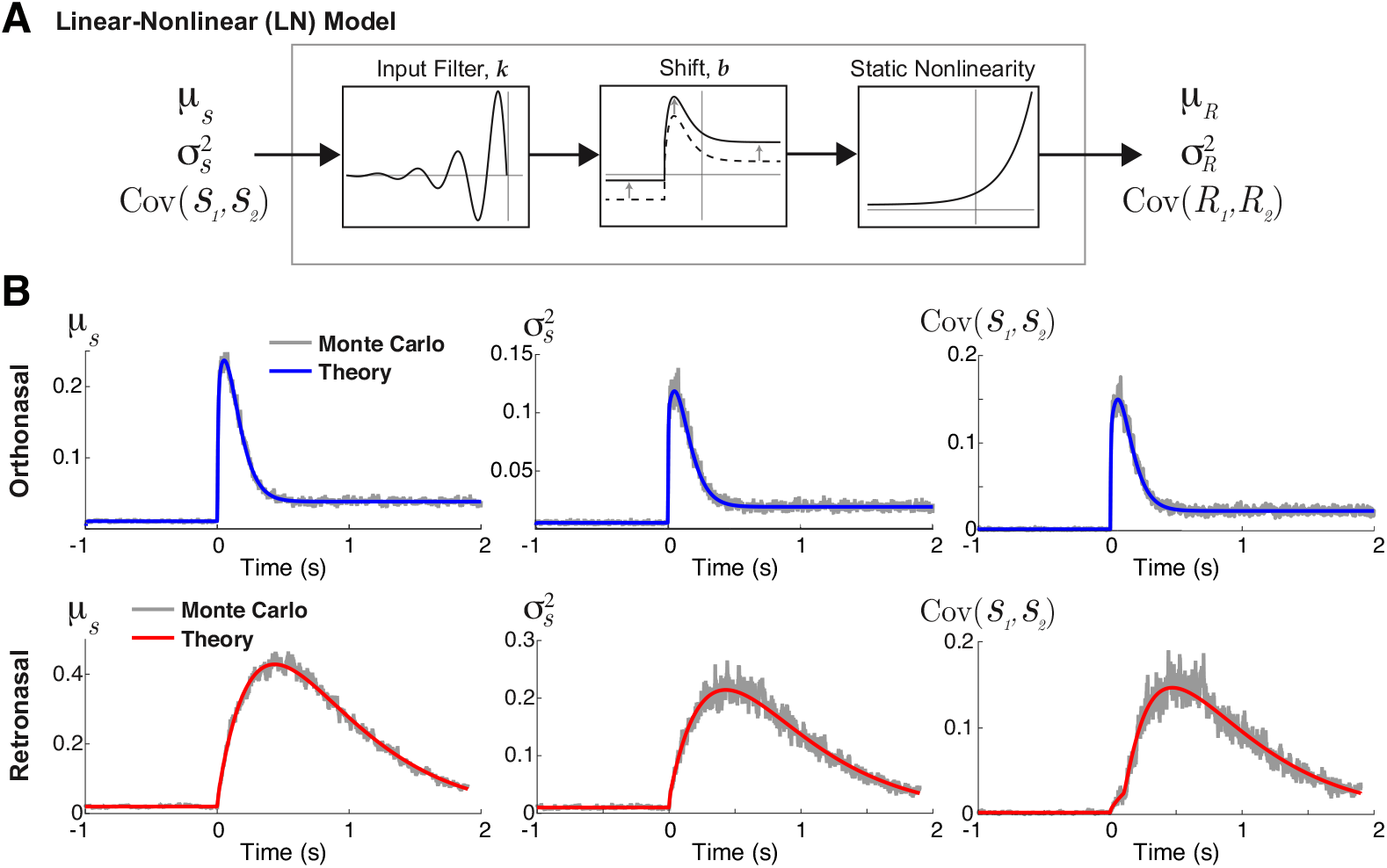
LN framework used to analyze OB transfer of input statistics. **A**) Schematic of the phenomenological linear-nonlinear (LN) model to approximate how the OB network transfers input statistics. **B)** The actual ortho (top row) and retro (bottom row) input synapses used in the biophysical OB model results in Figure 3. Comparisons of the Monte Carlo simulations (Eqs (10)–(11)) and theoretical calculations (Eqs (12), (17), (21) for respective columns) show smooth curve matches even for correlated time-varying (inhomogeneous) Poisson processes.

The LN model applies a temporal linear filter *k* (time-invariant) to the input *S*(*t*), and shifts the result by *b*, followed by a static non-linearity (exponential function) to produce an output *R*(*t*):

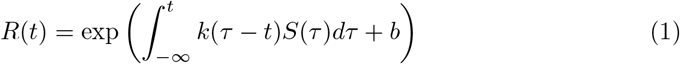

For our purposes, 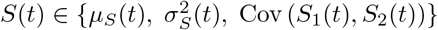 are the statistics of ORN input synapses, and 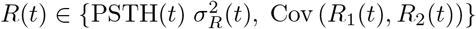 are the statistics of MC spiking response. For simplicity, we only consider:

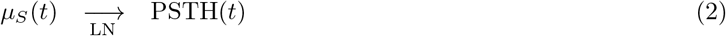

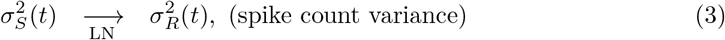

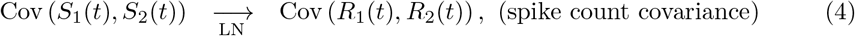

The LN model only accounts for specific input statistics transferred to their corresponding output statistics, without any mixing effects (e.g., 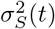 does not directly affect PSTH(*t*)). Although output statistics generally depend on all input statistics [28–30], we emphasize that our ad-hoc approach here is meant to better understand how the OB model operates and is not a principled alternative model.

For the inputs to the LN model, we use an exact theoretical calculation for *μ_S_* (*t*), 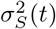, Cov(*S*_1_(*t*), *S*_2_(*t*)) rather than relying on Monte Carlo simulations. The ORN input synapses are driven by correlated time-varying inhomogenous Poisson processes yet we are still able to calculate the first and second order statistics of the ORN inputs in the limit of infinite number of realizations; detailed in **Materials and methods: Calculating time-varying ORN input synapses**, Eqs (12), (17), (21). A comparison of Monte Carlo simulations of the actual ORN inputs used in our OB model results (Eqs (10)–(11)) to the theoretical calculation (Eqs (12), (17), (21)) is shown in Fig 4B. We clearly see that the calculations (labeled ‘Theory’) matches all three ORN input statistics with smooth curves, properly accounting for both time-varying ORN input and time-varying input correlation. These calculations do not depend on any asymptotic assumptions; see Fig S2 in **S1. Supplementary Material** for more examples.

The fits of the LN model to the MC spike statistics (PSTH, variance, and covariance) are shown in Fig 5A. The LN model is a decent approximation to the output statistics obtained from the biophysical OB model for both ortho and retro stimuli. The resulting temporal filters *k*(*t*) in Fig 5B succinctly show how the various input statistics are filtered in time by the biophysical OB network model. For all 3 spike statistics, retro input statistics are filtered with larger absolute values (both positive and negative) than ortho, suggesting that the OB network is more sensitive to fluctuations with retro input. The resulting *b* values are listed in Table 1; the *b* represent an absolute shift independent of the temporal dynamics. They are similar for ortho and retro for all statistics except spike count covariance. Although *b* is important for the resulting LN curves (dot-black in Fig 5A), it is not a part of the temporal processing of ORN inputs.

**Fig 5.**
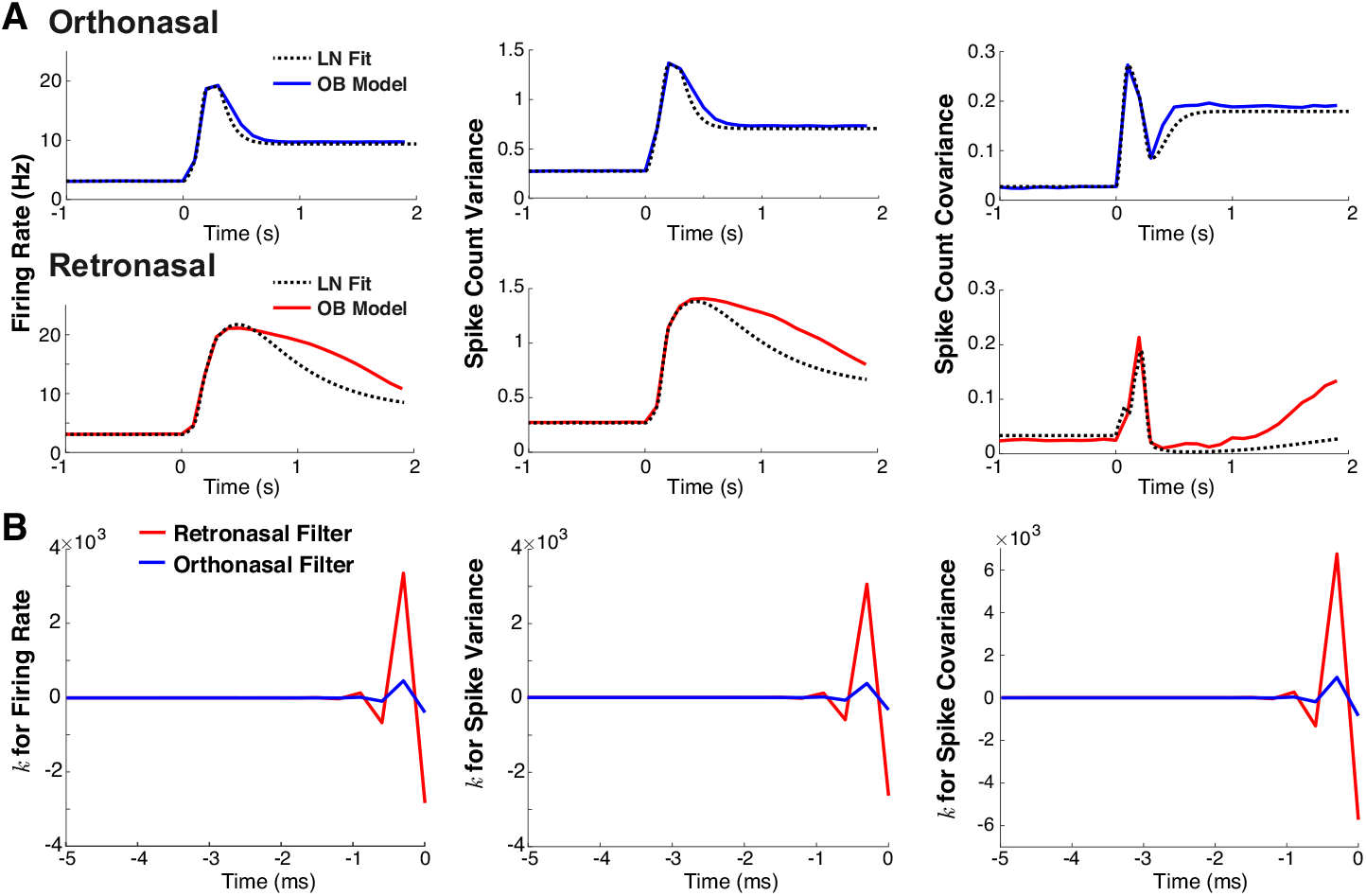
LN model shows that retronasal input results in temporal filters with larger magnitudes. **A**) Comparison of LN model output (dashed black curves) to OB network model output statistics for ortho (solid blue curves in top panels) and retro (solid red curves in bottom panels) stimulus with onset at *t* = 0 s. The LN output qualitatively captures OB model output statistics. **B**) Temporal filters *k*(*t*) in LN model for ortho (in blue) and retro (in red) stimulus over time (−5 ms≤ *t* ≤0 ms). Temporal filters for retro have larger positive and negative values than with ortho.

**Table 1.** Parameter *b* for LN model fits to MC spiking statistics in Figure 5.

|  | PSTH | Variance | Covariance |
| --- | --- | --- | --- |
| Orthonasal | 2.10 | -0.50 | -1.56 |
| Retronasal | 1.94 | -0.55 | -3.02 |

The parameter *b* for the LN model fits (Eq (1)) between orthonasal and retronasal are similar for a given statistic, except for spike count covariance.

### ORN input signatures for ortho/retro

Despite retro eliciting larger firing rates than ortho, the spike count covariance (as well as correlation and Fano factor) with retro stimulation is smaller than with ortho. It has long been known theoretically and experimentally that in uncoupled cells, the spike correlation increases with firing rate (at least with moderate to larger window sizes) [31], in contrast with our data. In coupled (recurrent) networks, the change in correlation with firing rate is complicated and depends on numerous factors [10–13]. Thus, the components of ORN inputs that result in these differences (higher firing and less covariance for retro than with ortho) in the same OB network are not obvious.

So we use our computational framework to uncover the important feature(s) of ORN input that: i) results in MC spike statistics consistent with our data trends, and ii) filters with larger absolute values with retro than with ortho input. Here we disregard the biological differences in ortho and retro ORN inputs to consider 3 core attributes of ORN inputs that influence how the OB model operates:

- Temporal (faster increase and decay, or slower increase and decay; see Fig 6A, left)
- Amplitude (low or high, see Fig 6A, left)
- Input correlation (low or high, black and gray curves respectively, in Fig 6A, right)

We consider a total of 8 different ORN input profiles consisting of various combinations of amplitude, input correlation, temporal profiles. The LN model fit to the OB model (i.e., MC spike statistics) for these 8 different ORN input profiles are all similar, well approximating how the OB coupled network transfers input statistics (see Fig 5A and Fig S3 in **S1. Supplementary Material**). Figure 6B clearly shows that the slower increase and decay in input rate (red and pink) consistently results in temporal filters *k*(*t*) with larger absolute values than with faster increase/decay (light blue and blue). This increased amplification in filter values is consistent with all 3 statistics, and is observed with all variations of amplitude and input correlation. Thus, the OB network consistently has temporal filters with larger absolute values when the input profile is slower (i.e., retronasal-like). The resulting LN model *b* values are listed in Table 2 for reference, although these values represent an absolute scaling independent of the temporal dynamics.

**Fig 6.**
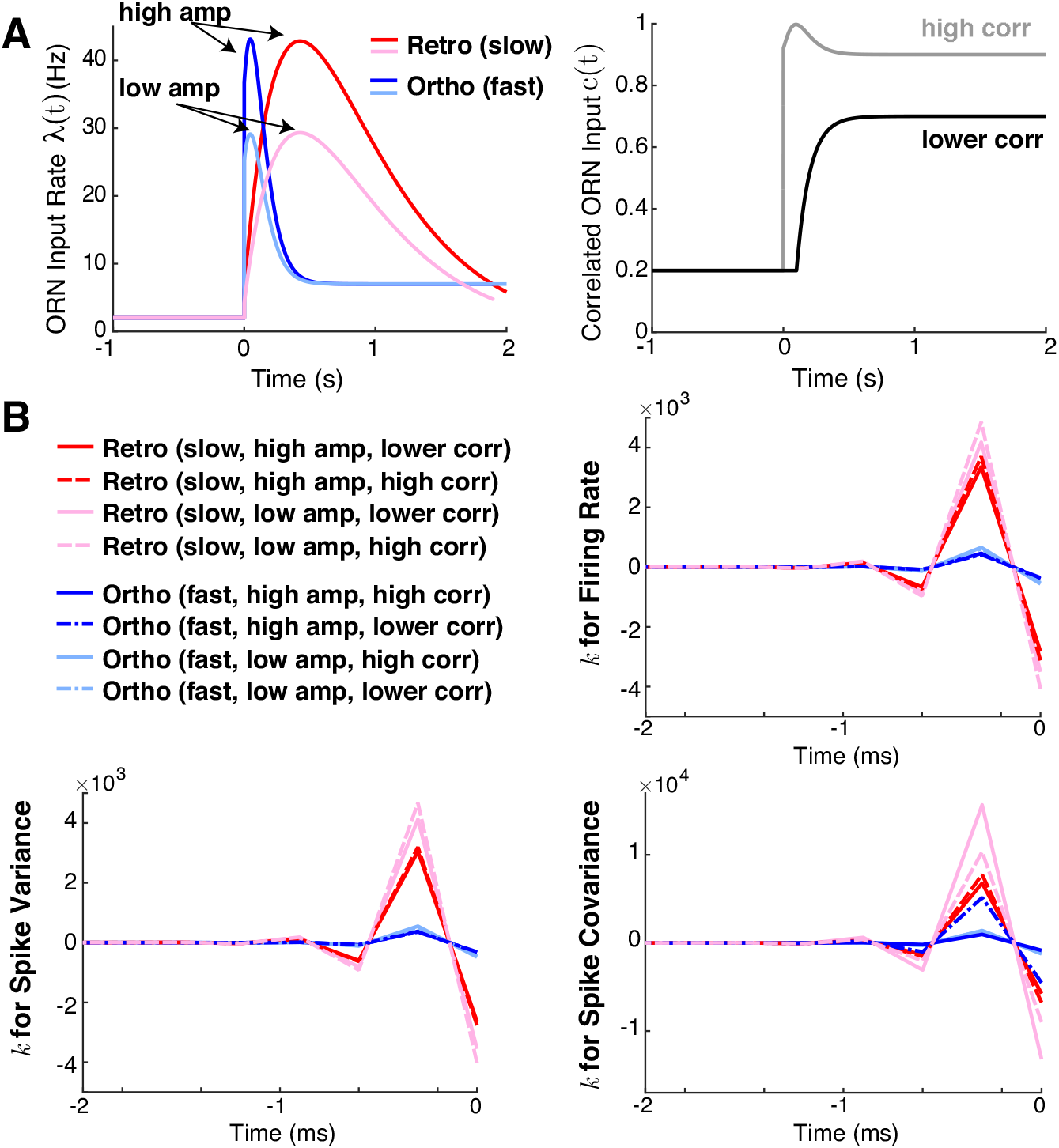
Temporal profile is crucial for larger magnitude filters. **A)** Different combinations of input rates (left) including slower increase and decay (retro-like) and faster increase and decay (ortho-like) as well as high and low amplitude as labelled. Two different input correlations (right), with high correlation in gray, and lower correlation in black. **B**) Resulting linear filters *k*(*t*) have consistently larger absolute values when temporal profile of ORN inputs is slower (red and pink), compared to faster (dark and light blue).

**Fig 7.**
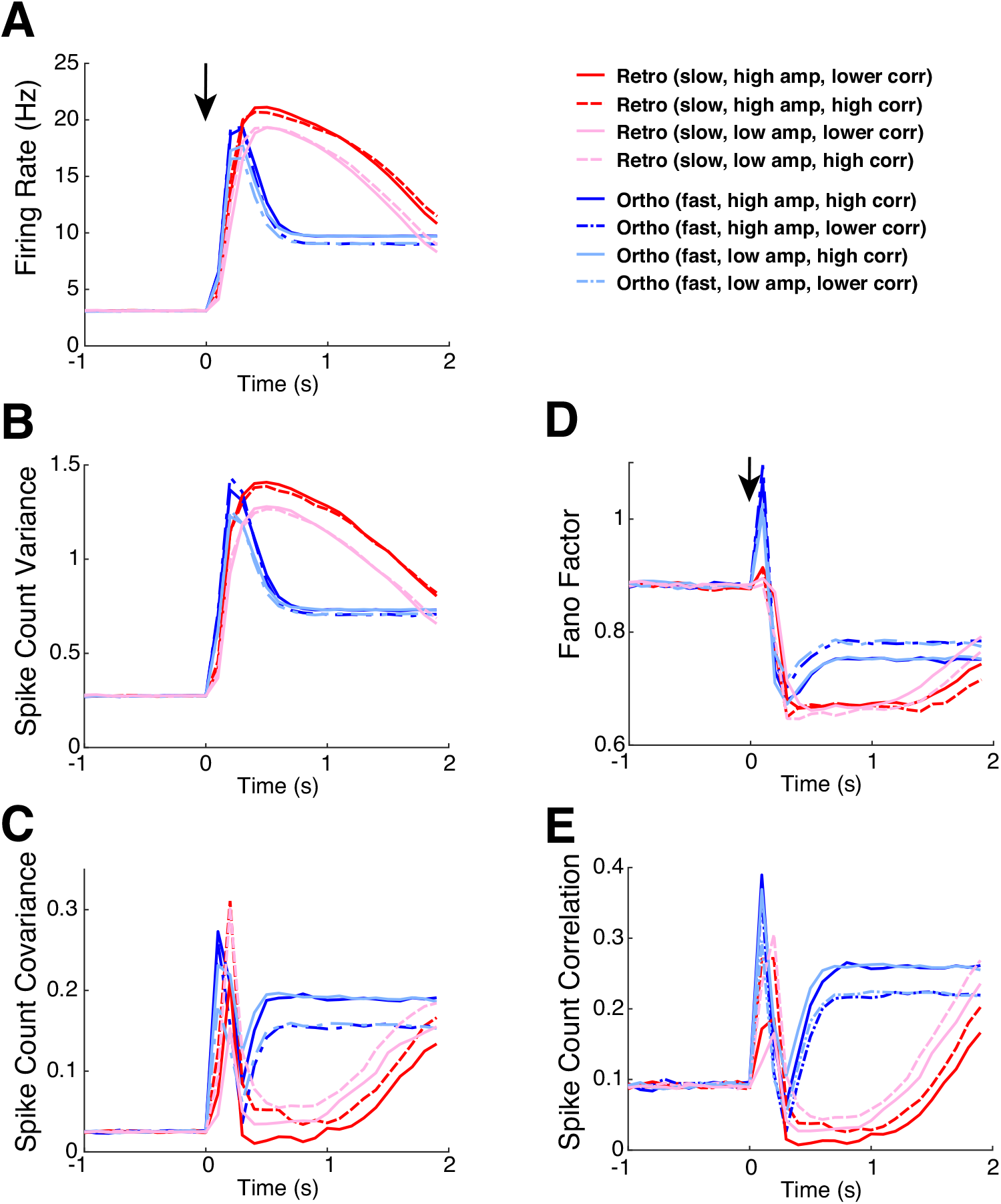
Comparison of all 8 OB model results. The 8 different OB model results are from varying temporal profile, amplitude height, and input correlation (2 ways each, see Fig 6A). Different temporal profiles is key for both having different model spike statistics **and** for best matching qualitative differences in our data (see Fig. 3). **A**) Firing rate in Hz (left) is slightly lower with low input rate amplitude, but no significant differences with different input correlations. **B**) Spike count variance, similar to firing rate, has only slightly lower values with low input rate amplitude. **C**) Spike count covariance is lower with lower input correlation for all of ortho evoked state (not surprisingly). However, retro (fast) with lower amplitude steadily increases above higher amplitude after about 1 s in the evoked state. **D**) Fano Factor model results only change modestly. **E**) Pearson’s spike count correlation, similar to spike count covariance, is lower with lower input correlation and similarly for retro (fast), there is an increase with higher input correlation.

**Table 2.** Parameter *b* for LN model fits to MC spiking statistics in Figure 6.

| Temporal | Amplitude | ORN correlation | PSTH | Variance | Covariance |
| --- | --- | --- | --- | --- | --- |
| Fast (ortho) | **High** | **High** | **2.10** | **-0.50** | **-1.56** |
|  | High | Low | 1.99 | -0.56 | -1.35 |
|  | Low | High | 2.03 | -0.55 | -1.62 |
|  | Low | Low | 1.97 | -0.61 | -3.42 |
| Slow (retro) | High | High | 2.04 | -0.50 | -1.11 |
|  | **High** | **Low** | **1.94** | **-0.55** | **-3.02** |
|  | Low | High | 1.70 | -0.72 | -1.36 |
|  | Low | Low | 1.59 | -0.81 | -3.18 |

The parameter *b* for the LN model fits (Eq (1)) of the various parameters for temporal profile, amplitude, and input correlation. Amplitude and ORN input correlation profiles as defined for Figs 3-5 and associated values previously listed in Table 1 are noted in bold.

Figure 7 shows all 8 OB model results for each spike statistic. For all first and second order statistics, including scaled measures of variability, the most distinct attribute that distinguishes our model results is the temporal profile of input. Importantly, the temporal profile is the key attribute to best capture the differences in ortho and retro our experimental data (see Fig. 3). The slow increase and decay in input rate consistently results in retro-like spiking trends while the fast increase and decay in input rate results in ortho-like spiking trends. Thus, our models show that the temporal profile is a signature of retro and ortho stimulation, and emphasizes the critical role of ORN inputs for shaping how the same OB network modulates ortho and retro stimuli.

## Discussion

We have investigated how odors processed via the orthonasal and retronasal routes result in different OB spike statistics, analyzing in detail how ORN inputs transfer to MC spike outputs. Motivated by our *in vivo* rat recordings that show significant differences in first and second order spiking statistics of MC, we developed a realistic OB network model to investigate the dynamics of stimulus-evoked spike statistic modulation (higher firing and lower covariance/correlation with retro than with ortho). Our OB model balances biophysical attributes [15, 16] with computational efficiency. The OB model is able to capture trends in our data with both ortho and retro stimulation, and should be useful for future studies of OB. We successfully used the biophysical OB model, paired with a phenomenological LN model, to analyze how different ORN inputs lead to different dynamic transfer of input statistics. We also showed that the temporal profile of ORN inputs is a key determinant of ortho versus retro input via model matching our data. The output spike statistics are crucial because the OB relays odor information to higher cortical regions, and thus our work may have implications for odor processing with different modes of olfaction.

To the best of our knowledge, our experiments detail the differences in MC spiking with ortho and retro stimuli for the first time. However, the work of Scott et al. [5] is related; they used 4 electrodes to record OB spiking activity in the superficial layers of OB in rats. Their results are difficult to directly compare to ours as they focus on superficial OB in the epithelium rather than the mitral cell layer, but at least the trial-averaged firing rates in their data appear to be consistent with our data. Moreover, our multi-electrode array recordings enable us to consider trial-to-trial covariance of spiking.

The key attribute(s) of ORN inputs that can result in different ortho and retro statistics consistent with our data are not obvious. Indeed, retro stimulation resulted in larger firing rates than ortho, and the spike count covariance (as well as correlation and Fano factor) with retro stimulation is smaller than with ortho, in contrast to uncoupled cells where correlation increases with firing rate [31]. Using various models, we were able to consider how three components of ORN inputs (temporal profile, amplitude, and input correlation) result in different OB dynamics with regards to transferring input statistics to outputs. Prior experiments [1–3] have shown these input components can differ with ortho and retro inputs. We found that the temporal profile (fast versus slow) plays a critical role for both capturing our data and for shaping the transfer of inputs to outputs, i.e., retro inputs consistently resulted in larger temporal filter values, so the OB network is more sensitive to fluctuations of retro input statistics than ortho. The slower input rate (rise and decay) is a key signature of retronasal stimulation [2, 3, 5] to capture the trends in our data with retro stimulation, while faster rise and decay is similarly a key signature of orthonasal stimulus.

The temporal differences between ortho versus retro have previously been thought to play a role in distinguishing ortho/retro stimulation at the ORN [1–3, 5, 6], but whether this carried over to the OB and if this held at the level of spiking was unknown. Here we demonstrate the importance of different temporal input to OB for ortho versus retro.

We used an ad-hoc LN model framework because many of biological complexities are removed yet important features are retained. That is, neurons are known to linearly filter inputs, i.e., finding linear filters of neurons is not new [27], and they are related to the spike-triggered average [32], and spike generation is inherently nonlinear. Thus, LN-type models have been used in many contexts, and often to circumvent biophysical modeling, most notably with generalized linear models [33, 34] (also see [26]) where various filters and model components are fit to data using maximum likelihood. Connecting the large gap between biophysical models and LN models is daunting, but see Ostojic and Brunel [25] who relate stochastic integrate-and-fire type models to LN. Our approach here is much simpler than the aforementioned works because we simply wanted to assess how a particular statistic (mean, variance or covariance) mapped via the OB network in a simple and transparent manner.

With a combination of experiments and different scales of neural network modeling, we provide a basis for understanding how differences in OB spiking statistics arise with these 2 natural modes of olfaction. More generally, our model framework provides a road map for how to analyze attributes responsible for different OB spiking when driven by differences in ORN inputs.

## Materials and methods

See https://github.com/michellecraft64/OB for MATLAB code implementing the single-compartment biophysical model, the equations for synaptic input statistics, and the linear-nonlinear (LN) model.

### Single-compartment biophysical OB model

Models of all three cell types (MC, PGC, GC) are based on models developed by the Cleland Lab [15, 16]. We consider two glomeruli each with a representative MC, PGC, GC (see Fig 2B). Each cell is a conductance-based model with intrinsic ionic currents. The voltage responses of all three cell types, measured in experiments and in a multi-compartment model [15, 16], are generally captured in our single-compartmental model, see Fig 2A. Here we describe all of the pertinent model details thoroughly; for other extraneous details and implementation, please refer to provided code on GitHub.

### Individual cell model

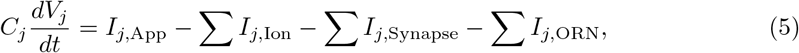

The voltages of all model cells are governed by a Hodgkin-Huxley type current balance equation (Eq (5) above for the *j*^th^ cell) consisting of voltage (*V*), membrane capacitance (*C*), applied current (*I*_App_), ionic currents (*I*_Ion_), synaptic currents (*I*_Synapse_), and ORN inputs (*I*_ORN_); see Table 3 and 4 for units and numerical values, respectively. For our modeling purposes, the ionic currents and the ORN inputs are modified from [15, 16] and described below.

**Table 3.**
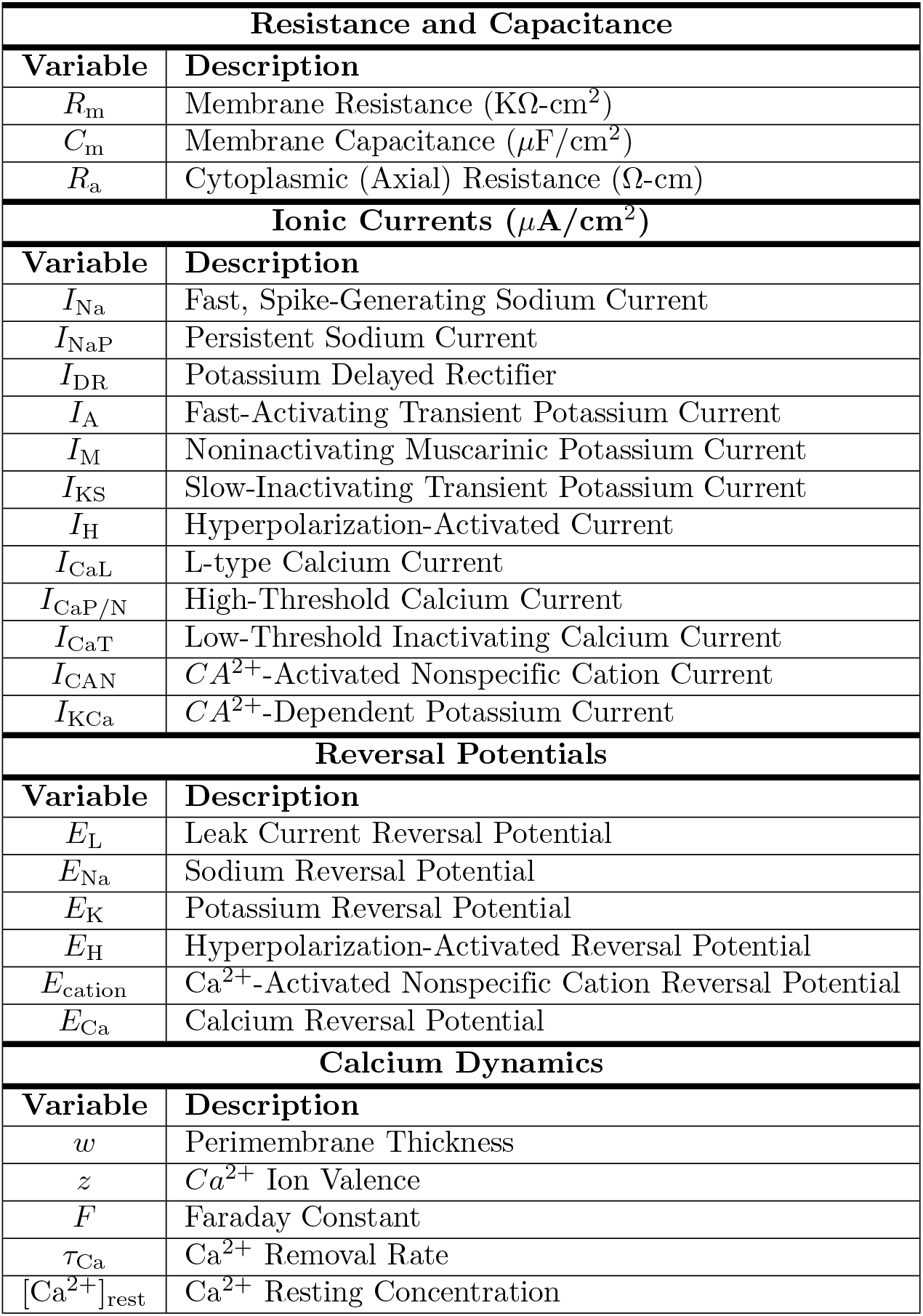
Description of model parameters.

**Table 4.** Parameter values for each cell type. Each of these values are the same as defined by [16] with the exception of maximal conductance values which are the sum of all cell compartments (soma, dendrite, axon, etc.) as defined by [16]. Additionally, any conductance value denoted by implies that this ionic current is not included in the associated cell.

| Resistance and Capacitance |  |  |  |
| --- | --- | --- | --- |
| Variable | MC Value | GC Value | PGC Value |
| $R_m$ | 30 | 30 | 20 |
| $C_m$ | 1.2 | 2.0 | 1.2 |
| $R_a$ | 70 | 70 | 80 |
| Maximal Conductance (mS/cm<sup>2</sup>) |  |  |  |
| Variable | MC Value | GC Value | PGC Value |
| $g_{Na}$ | 120 | 70 | 70 |
| $g_{NaP}$ | 0.42 | — | — |
| $g_{DR}$ | 70 | 25 | 25 |
| $g_A$ | 10 | 80 | 40 |
| $g_M$ | — | 0.5 | 1.0 |
| $g_{KS}$ | 84 | — | — |
| $g_H$ | — | — | 0.2 |
| $g_{CaL}$ | 0.85 | — | — |
| $g_{CaP/N}$ | — | 0.2 | 1.0 |
| $g_{CaT}$ | — | 0.1 | 3.0 |
| $g_{CAN}$ | — | 1.0 | — |
| $g_{KCa}$ | 5 | 0.5 | 2.0 |
| Reversal Potentials (mV) |  |  |  |
| Variable | MC Value | GC Value | PGC Value |
| $E_L$ | -60 | -60 | -65 |
| $E_{Na}$ | 45 | 45 | 45 |
| $E_K$ | -80 | -80 | -80 |
| $E_H$ | 0 | 0 | 0 |
| $E_{cation}$ | 10 | 10 | 10 |
| Calcium Dynamics |  |  |  |
| Variable | MC Value | GC Value | PGC Value |
| $w$ | 1 $\mu m$ | 0.2 $\mu m$ | 0.2 $\mu m$ |
| $z$ | 2 | 2 | 2 |
| $\tau_{Ca}$ | 10 ms | 800 ms | 800 ms |
| $[Ca^{2+}]_{rest}$ | 0.05 $\mu mol/l$ | 0.05 $\mu mol/l$ | 0.05 $\mu mol/l$ |

### Ionic currents

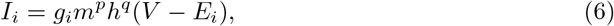

The ionic currents are defined by Eq (6) above (for specific ion type *i*) and account for maximal conductance (*g*), activation variable (*m*) with exponent (*p*), inactivation variable (*h*) with exponent (*q*), time-varying voltage (*V* assumed to be isopotential), and reversal potential (*E_i_*). All parameters and function for intrinsic ionic currents and their gating variables are the same as in [15, 16] with the exception of maximal conductance. We chose to condense the model as defined in [15, 16] by collapsing all compartments to a single-compartment, and we set the maximal conductance as the sum of all maximal conductance values (e.g., in PGC, *I*_Na_ has maximal conductance *g*_Na_ = 70 mS/cm^2^ because [15] set *g*_Na_ = 50 mS/cm^2^ in the soma and *g*_Na_ = 20 mS/cm^2^ in the spine). All summed maximal conductance values used are listed for reference in Table 4. The calcium dynamics used to define the calcium-related ionic currents are the same as in [15, 16].

### Synaptic currents

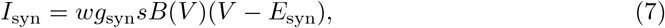

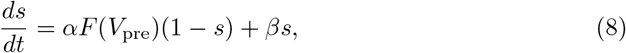

Eqs (7) and (8) are the equations for the synaptic variables, where all presynaptic GCs and PGCs provide GABA_A_ inputs, and all presynaptic MCs provide both AMPA and NMDA inputs. *B*(*V*) in Eq (7) is the NMDA-specific magnesium block function (*B*(*V*) = 1 for all other synapses), and *s*(*t*) is the fraction of open synaptic channels. The channel opening rate constants (*α* and *β*) are normalized sigmoidal function of presynaptic membrane potential (*F* (*V*_pre_) in Eq (8)), the same as in [15, 16]. We also define the conductance parameter (*g*_syn_) and reversal potentials (*E*_syn_) as [15, 16] have, with *g*_GABA_ = 1.5 nS for GC→MC synapses, *g*_GABA_ = 2 nS for PGC→MC synapses, *g*_AMPA_ = 2 nS and *g*_NMDA_ = 1 nS for both MC→PGC and MC→GC synapses; *E*_syn_ = 0 mV for AMPA and NMDA currents, and *E*_syn_ = −80 mV for GABA_A_ currents.

In order to capture different effects of coupling, we include coupling strengths *w*. The synaptic coupling strengths are fixed and set to: *w_M←G_* = 3 (independent inhibition), *w_M←Gc_* = 0.3 (common inhibition to MC), *w_G←M_* = 1 (same for both AMPA, NMDA), *w_Gc←M_* = 0.5 (inhibition to common GC), *w_P ←M_* = 1 and *w_M←P_* = 2 (same for both AMPA, NMDA). These coupling strengths were established based on preliminary results by Ly et. al [35] who use a related biophysical OB network model to evaluate regions of parameter spaces that provide best model fits to our experimental data. Similar to as seen in Ly et. al [35] (see their Figs 2 and 3) we define independent inhibition defined for GC to be greater than excitation from MC and common GC inhibition to be less than both GC inhibition and MC excitation (i.e., *w_Gc←M_* ≤ *w_G←M_* ≤ *w_M←G_*).

### ORN input

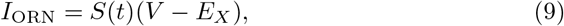

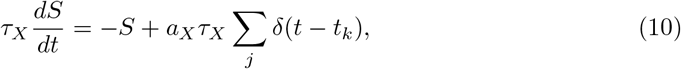

The ORN inputs for each cell consist of both excitatory and inhibitory inputs as specified in Eqs (9) and (10) where *X* ∈ {*E, I*}. The reversal potential value (*E_X_*) is much larger for excitatory inputs and smaller for inhibitory. The function 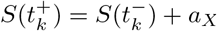 accounts for the random times (*t_k_*) when *S* instantaneously increases by *a_X_*. The random times, *t_k_*, are governed by an inhomogeneous Poisson process with rate *λ_X_* (*t*). This aligns with experimental evidence that ORN spiking is Poisson-like in the spontaneous state [9]. Thus, we extend the notion that ORN spiking would be Poisson-like in the evoked state with increased rate *λ_X_* (*t*) varying in time. Finally, we set the synaptic rise and decay time constants (*τ_X_*) to be 5.5 ms for PGCs and GCs, 10 ms for MCs, as in [15, 16].

In order to account for odor input response (i.e, spontaneous to evoked states), as well as differentiating between ortho- vs. retronasal odor input, we modulate the time-varying Poisson input rate (*λ_X_* (*t*)). The ortho- vs. retronasal odor input rates, *λ_O/R_*(*t*), were constrained such that *λ_O_*(*t*) increases faster and more abruptly than *λ_R_*(*t*) with odor, and *λ_R_*(*t*) decays slower than *λ_O_*(*t*) based on ORN imaging studies [2, 3]. The time-varying *λ_O/R_*(*t*) for ortho- and retronasal stimulus can be seen plotted in blue and red, respectively, in Fig 2Ci. Inputs consist of both excitatory synapses (with rate *λ_O/R_*(*t*)) and inhibitory synapses (with rate 0.75*λ_O/R_*(*t*)) to capture other unmodeled inhibitory effects. For specific algebraic formula of *λ_R/O_*(*t*), please refer to code found in listed GitHub link.

The ORN input rates can be pairwise correlated, which is achieved by the parameter *c_j,k_* ∈ [0, 1], for cells *j* and *k* detailed by Eq (11) below:

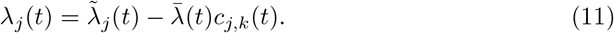

where 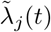 and 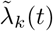 are the individually defined ORN input rates for cells *j* and *k*, and 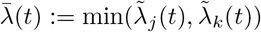.

We set the correlation (*c_j,k_*) between the following cell pairs: MC and PGC pair within a glomerulus that have inputs from the same ORN cells (*c_j,k_* = 0.3); two MCs as they are known to have correlated ORN input [36] (*c_j,k_*(*t*) time-varying as in Fig 2Cii); and between all 3 GCs because they are known to synchronize with common input [16, 37] (*c_j,k_* = 0.3). All other pairs of cells have no background input correlation. Specifically, input correlation for the 2 MCs (*c_j,k_*(*t*)) varied in time to mimic increased correlation of glomeruli activity with odor onset and additionally account for ortho vs. retronasal odor input (*c_O/R_*(*t*)). As seen in Fig 2Cii, input correlation for the 2 MCs are constrained such that *c_R_*(*t*) < *c_O_*(*t*). This constraint is based on prior imaging studies showing smaller spatial maps of retronasal response, and more specifically that retronasal spatial maps are subsets of orthonasal [2, 3]. For specific algebraic formula of *c_R/O_*(*t*), please refer to code found in listed GitHub link.

### Calculating time-varying ORN input statistics of synapses

Here we describe a method to capture the effects of ORN input statistics of synapses to the biophysical OB model, in the limit of infinite realizations. These methods are very useful as inputs for the LN model, without which one would have to use averages from Monte Carlo simulations that contain deviations from finite size effects. Taking the expected value of Eq (10) results in an equation for the average of *S*(*t*), *μ_S_* (*t*):

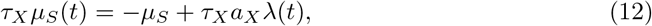

To derive the equation for variance 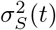, we multiply Eq (10) by itself. We can equivalently rewrite Eq (10) as an integral:

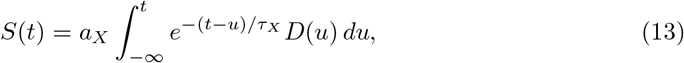

where 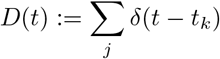. So

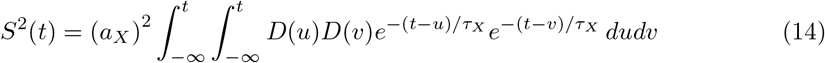

Recall that ***E***[*D*(*u*)*D*(*v*)] = *λ*(*v*)*δ*(*u* − *v*) + *λ*(*u*)*λ*(*v*), so we have:

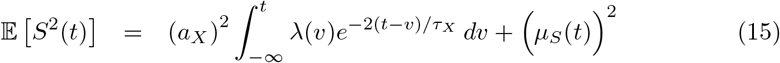

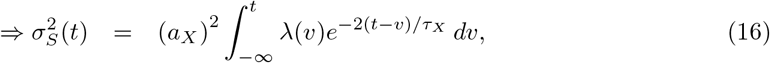

Equivalently, 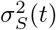 satisfies the ODE:

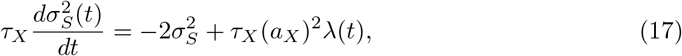

Similarly for *S_j_*(*t*)*S_k_*(*t*) correlated synapses, we have:

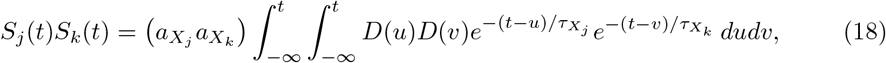

By our model construction 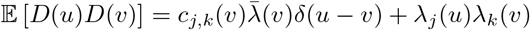, where 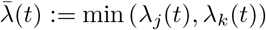, so we have:

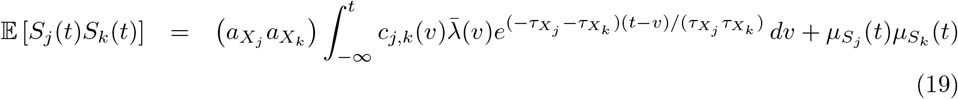

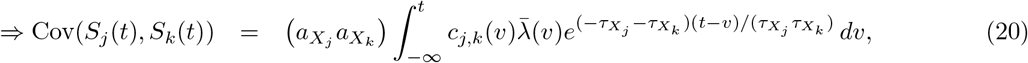

Cov(*S_j_*(*t*), S_k_(*t*)) equivalently satisfies the ODE:

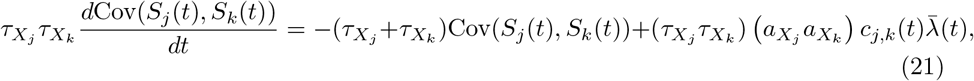

The calculations for the dynamic (time-varying) synapse statistics are important for capturing realistic statistics because a steady-state approximation can be very inaccurate, especially when the time-varying correlation and ORN spiking rate change quickly relative to the time-scales (*τ_X_*). The quasi-steady-state approximation is:

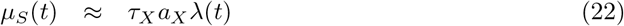

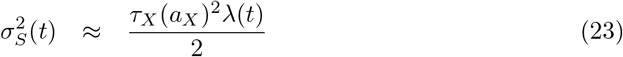

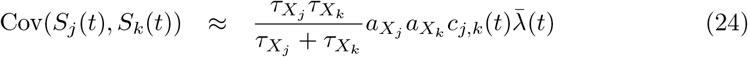

Figure S2 in **S1. Supplementary Material**shows several more examples demonstrating the accuracy of the calculations (Eqs (12), (17), (21)) and how inaccurate Eqs (22)–(24) can be.

### Linear-Nonlinear (LN) model

We use the above described ODEs (Eqs (12), (17), and (21)) to simplify the ORN input statistics calculations for use with the LN model framework. Previous work has implemented LN-type models as an alternative to biophysical spiking models with various conditions (see [25–27] and Discussion). Fig 4A illustrates a schematic of the LN model parameters, linear filter (*k*) and shift (*b*), that are used with the ORN input statistics 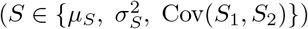 in order to calculate output spike statistics 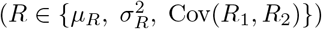. The LN model is summarized as:

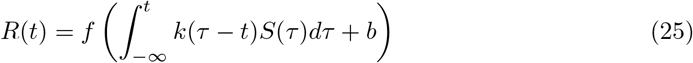

Where we define our function *f* as an exponential, and we can approximate the integral numerically as follows:

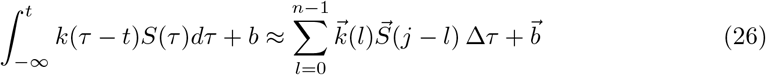

Where *n* denotes the number of time points included in the linear filter, and *j* denotes the points in time of input statistic 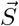 of size Lt. Then, we can rewrite Eq (26) in matrix vector form 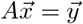 where: *A* is the Toeplitz matrix of size (Lt − *n* + 1) × *n* of our input statistic 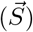 with an additional row of value one to account for shift; 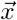 is our linear filter 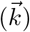 and shift 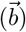; and 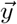 is our OB network firing rate statistic to which we fit our filter. Then, we solve for 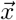 using Least Squares approximation by QR decomposition. The linear filter (*k*) converges to 0 by construction, therefore we truncate the filter at −0.1 s and set *k* = 0 for the remaining time −1 ≤ *t* < −0.1. Then, the output firing rate statistic (*R*) is calculated as follows:

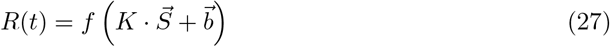

Where *K* denotes the convolutional matrix constructed from the truncated linear filter *k*.

### Electrophysiological recordings

We decided to use recordings from a single rat, with recordings from 3 sessions. We took this conservative approach to control differences in nasal cavity structure that can vary across rats [38, 39], which may shape differences in ortho versus retro activity [2, 40]. See provided GitHub code for statistical summary of experimental data.

All procedures were carried out in accordance with the recommendations in the Guide for the Care and Use of Laboratory Animals of the National Institutes of Health and approved by University of Arkansas Institutional Animal Care and Use Committee (protocol #14049). Data were collected from 11 adult male rats (240-427 g; *Rattus Norvegicus*, Sprague-Dawley outbred, Harlan Laboratories, TX, USA) housed in an environment of controlled humidity (60%) and temperature (23°C) with 12h light-dark cycles. The experiments were performed in the light phase.

#### Surgical preparations

Anesthesia was induced with isoflurane inhalation and maintained with urethane (1.5 g/kg body weight (**bw**) dissolved in saline, intraperitoneal injection (**ip**)). Dexamethasone (2 mg/kg bw, ip) and atropine sulphate (0.4 mg/kg bw, ip) were administered before performing surgical procedures. Throughout surgery and electrophysiological recordings, core body temperature was maintained at 37°C with a thermostatically controlled heating pad. To isolate the effects of olfactory stimulation from breath-related effects, we performed a double tracheotomy surgery as described previously [2]. A Teflon tube (OD 2.1 mm, upper tracheotomy tube) was inserted 10mm into the nasopharynx through the rostral end of the tracheal cut. Another Teflon tube (OD 2.3 mm, lower tracheotomy tube) was inserted into the caudal end of the tracheal cut to allow breathing, with the breath bypassing the nasal cavity. Both tubes were fixed and sealed to the tissues using surgical thread. Local anesthetic (2% Lidocaine) was applied at all pressure points and incisions. Subsequently, a craniotomy was performed on the dorsal surface of the skull over the right olfactory bulb (2 mm × 2 mm, centered 8.5 mm rostral to bregma and 1.5mm lateral from midline).

#### Olfactory Stimulation

A Teflon tube was inserted into the right nostril and the left nostril was sealed by suturing. The upper tracheotomy tube inserted into the nasopharynx was used to deliver odor stimuli retronasally. Odorized air was delivered for 1 s in duration at 1 minute intervals. The odorant was Ethyl Butyrate (EB, saturated vapor). We note that the full experimental data set included additional odors, but here we consider only EB.

#### Electrophysiology

A 32-channel microelectrode array (MEA, A4×2tet, NeuroNexus, MI, USA) was inserted 400 *μ*m deep from dorsal surface of OB targeting tufted and mitral cell populations. The MEA probe consisted of 4 shanks (diameter: 15 *μ*m, inter-shank spacing: 200 *μ*m), each with eight iridium recording sites arranged in two tetrode groups near the shank tip (inter-tetrode spacing: 150 *μ*m, within tetrode spacing 25 *μ*m). Simultaneous with the OB recordings, we recorded from a second MEA placed in anterior piriform cortex. Voltage was measured with respect to an AgCl ground pellet placed in the saline-soaked gel foams covering the exposed brain surface around the inserted MEAs. Voltages were digitized with 30 kHz sample rate (Cereplex + Cerebus, Blackrock Microsystems, UT, USA). Recordings were band-pass filtered between 300 and 3000Hz and semiautomatic spike sorting was performed using Klustakwik software, which is well suited to the type of electrode arrays used here [41].

## Supporting Information

**S1. Supplementary Material. Other parts of the research that support the main results but are separated for a more streamlined exposition.**

## Acknowledgments

We thank the Southern Methodist University (SMU) Center for Research Computing for providing computational resources. This work was supported by the National Science Foundation (#IIS-1912338 Cheng Ly and Michelle Craft, #IIS-1912320 Andrea Barreiro, #IIS-1912352 Woodrow Shew and Shree Hari Gautam).

## Author Contributions

MC and CL developed the computational models. MC programmed and implemented the software. MC, AKB, WLS, CL conceptualized the project. SHG and WLS designed the experiments and collected the data. MC and CL drafted the original manuscript. MC, AKB, SHG, WLS, CL edited the manuscript. MC and CL developed the visualizations. CL supervised the project.

